# Improving antibody-mediated protection against HSV infection by eliminating interactions with the viral Fc receptor gE/gI

**DOI:** 10.1101/2024.11.20.624598

**Authors:** Matthew D. Slein, Iara M. Backes, Natasha S. Kelkar, Callaghan R. Garland, Urjeet S. Khanwalkar, Anton M. Sholukh, Christine M. Johnston, David A. Leib, Margaret E. Ackerman

## Abstract

Herpes simplex virus (HSV) encodes surface glycoproteins that are host defense evasion molecules, allowing the virus to escape immune clearance. In addition to their role in neuropathogenesis and cell-cell spread, glycoproteins E and I (gE/gI) form a viral Fc receptor (vFcR) for most subclasses and allotypes of human IgG and promote evasion of humoral immune responses. While monoclonal antibodies (mAbs) protect mice from neonatal HSV (nHSV) infections, the impact of the vFcR on mAb-mediated protection by binding to IgG is unknown. Using HSV-1 with intact and ablated gE-mediated IgG Fc binding, and Fc-engineered antibodies with modified ability to interact with gE/gI, we investigated the role of the vFcR in viral pathogenesis and mAb-mediated protection from nHSV. The gD-specific human mAb HSV8 modified to lack binding to gE exhibited enhanced neutralization and *in vivo* protection compared to its native IgG1 form. This improved protection by the engineered mAbs was dependent on the presence of the vFcR. Human IgG3 allotypes lacking vFcR binding also exhibited enhanced antiviral activity *in vivo*, suggesting that vaccines that robustly induce IgG3 responses could show enhanced protection. suggesting the value of vaccination strategies that robustly induce this subclass. Lastly, analysis of longitudinal responses to acute primary genital infection in humans raised the possibility that unlike most viruses, HSV may exhibited slow induction of IgG3. In summary, this study demonstrates that mAbs lacking the ability to interact with the vFcR can exhibit improved protection from HSV—offering new prospects for antibody-based interventions.

**One Sentence Summary:** The herpes simplex virus neutralizing antibody HSV8 demonstrates improved activity *in vitro* and *in vivo* when its IgG Fc domain lacks the ability to bind the viral Fc receptor glycoprotein E/I complex through either Fc engineering or natural human IgG3 allotypes.

## Introduction

Herpes simplex viruses are enveloped double-stranded DNA viruses that are ubiquitous in human populations primarily due to their unique life cycles and multiple strategies for evasion of the host immune system *(1)*. HSV-1 and HSV-2 primarily infect mucosal surfaces before traveling to the peripheral nervous system and infecting sensory ganglia *(2)*. The development of a successful vaccine has been elusive due to the ability of HSV to establish latency and its ability to evade innate and adaptive immune responses (*3, 4)*. In lieu of an efficacious vaccine, monoclonal antibodies (mAbs) may serve as potent antiviral therapies to augment the standard of care— especially for vulnerable populations such as immunocompromised individuals and neonates, who are at higher risk from severe primary and/or reactivated disease (*5, 6)*.

Pre-clinical studies utilizing mAbs that can neutralize and/or mediate effector functions have demonstrated great efficacy for HSV prevention *(7–12)*, motivating translation of mAbs to clinical trials for both preventative and therapeutic purposes in adults. However, these trials have yet to report success, and there are currently no approved mAb treatments. Consistent with this gap between pre-clinical and clinical prospects, epidemiological evidence is somewhat split. The protective role of antibodies is well-established in the setting of primary infection, both in neonates and in adults. Birthing parent HSV seropositivity greatly reduces the risk of infection in neonatal HSV acquisition (<3% vs. 50%) *(13–17)* in association with transplacental transfer of IgG to the fetus, and HSV-1 seropositivity in adults decreases the severity of HSV-2 infection *(18)*. Further, protection from HSV-1 infection afforded by a candidate vaccine based on HSV-2 antigens was correlated with neutralization titers against HSV-1 (*19, 20)*. Other studies support the notion that antigen specificity, antiviral functions such as reducing cell-cell spread, antibody and IgG receptor polymorphisms, as well as humoral immune response deficiencies, could contribute to the severity and/or frequency of HSV reactivation *(21–24)*. On the other hand, the presence of endogenous binding and neutralizing Abs is usually insufficient to prevent reactivation (*21, 25)*. Such gaps in the protective capacity of antibodies raise questions as to how to improve the activity of mAb therapeutics or vaccines.

To this end, one hurdle in the development of an efficacious vaccine or mAb therapeutic is viral evasion of humoral immunity through glycoproteins E and I (gE/gI). The gE/gI heterodimer is expressed on the surface of HSV and infected cells (*26, 27)* and is critical for cell-cell spread, establishment of latency *(28–30)*, and neuropathogenesis (*31, 32)*. Beyond these roles, gE/gI also binds to the Fc region of IgG, allowing the virus to evade humoral immunity by functioning as a viral Fc receptor (vFcR) *(33–36)*. The gE/gI complex binds to human IgG1, IgG2, and IgG4 subclasses, but not all IgG3 allotypes, and can protect virally infected cells from antibody-dependent cellular cytotoxicity (ADCC) and antibody-dependent complement activity *(37–40)*. Additionally, the vFcR aids in humoral immune evasion through a mechanism known as antibody-bipolar bridging (ABB), in which antigen-bound antibody is simultaneously bound by gE/gI on the Fc region (*41, 42)*. This ternary complex can then be endocytosed and through the pH-dependent binding of gE to Fc, IgG is released in acidifying endosomes and subsequently degraded *(42)*. The vFcR may traffic back to the surface, potentially enabling continuous degradation of HSV-specific antibodies. The disparity of vFcR binding among IgG subclasses suggests that engineering a better antibody is possible and that improving our understanding of gE/gI mediated humoral immune evasion has the potential to contribute to antibody-based interventions.

Of the different clinical indications of HSV infections in humans, nHSV is perhaps the most amenable to antibody (Ab) therapeutics and vaccination strategies. While cases are rare, nHSV frequently results in severe life-long morbidity, and neonates contend with high mortality rates even with antiviral therapy *(43–45)*. Our work has previously demonstrated that mAbs can protect neonatal mice from HSV-mediated mortality *(7)* through both neutralization and effector functions *(8)*. Beyond clinical relevance, neonatal mice provide a useful model system to decipher Ab-virus interactions to enhance understanding of how Abs may protect against HSV infections more broadly.

Here, using an Fc engineering approach for IgG1 and naturally-evading IgG3 mAbs, as well as mutated virus, we investigate the role of vFcR activity in mAb-mediated protection. We find that the vFcR reduces mAb efficacy, and that optimizing antibody antiviral activity through Fc- and subclass-engineering can significantly improve the efficacy of HSV8 in the prevention and treatment of viral infection.

## Results

### gD-specific antibodies with Fc mutations extending half-life provide improved protection independent of mAb pharmacokinetics

HSV8 *(9)*, a potently neutralizing IgG1 antibody targeting the viral entry mediator gD, protects adult and neonatal mice from HSV-1-mediated mortality (*8, 9)* and has been evaluated for safety in a phase I clinical trial (NCT02579083) *(46)*. With an eye towards clinical translation, we evaluated the effect of Fc mutations that extend serum half-life *(47)* via interactions with the human neonatal Fc receptor (FcRn) and which are becoming standard in mAb therapies for infectious diseases. We evaluated variants of HSV8 IgG1 with three common human half-life extension mutations: M252Y/S254T/T256E (YTE) *(48)*, M428L/N434S (LS or Xtend) *(49)*, and M428L/N434A (LA) *(50)*. In humans, including neonates, these mutations have been shown to increase antibody half-life from ∼21 days to upwards of 100 days for some antibodies *(47)* — motivating investigation of the impact of these Fc mutations on mAb-mediated protection in the mouse model of nHSV-1 infection. To evaluate whether the efficacy of HSV8 was compromised by the inclusion of the half-life extending Fc mutations, 2-day old C57BL6/J (B6) mice were injected intraperitoneally (i.p.) with either 40 or 10 µg of mAb and then immediately challenged with 10^4^ plaque forming units (PFU) of HSV-1 st17 intranasally (i.n.).

Each HSV8 variant robustly protected mice from a lethal HSV-1 infection at both high (40 µg) (**Figure 1A**) and low (10 µg) (**Figure 1B**) doses — providing statistically significant protection (p<0.0001) as compared to an isotype control (VRC01) *(51)*. While similar levels of protection for unmodified (WT) and human half-life extension mutants (LS, LA, and YTE) were observed at the 40 µg dose, we observed striking differences at the 10 µg dose. HSV8 LA and HSV8 LS provided significantly better protection than the WT IgG1, whereas 10 µg of the WT HSV8 IgG1 and HSV8 YTE protected only about 50% of pups (**Figure 1B**). Weights among surviving pups were similar (**Figure S1**). The variable impact of human half-life extension mutations on antibody effector function provided a potential explanation of variable protection observed: YTE mutations extend half-life at the cost of reduced FcγR affinity and ability to mediate effector functions *(52)*, which are known to contribute to protection *(8)*. Given the short time frame of this challenge model, and prior reports that these mutations do not improve antibody pharmacokinetics (PK) and half-life in mice, we sought an explanation for the improved protection seen at the 10 µg dose.

**Figure 1:**
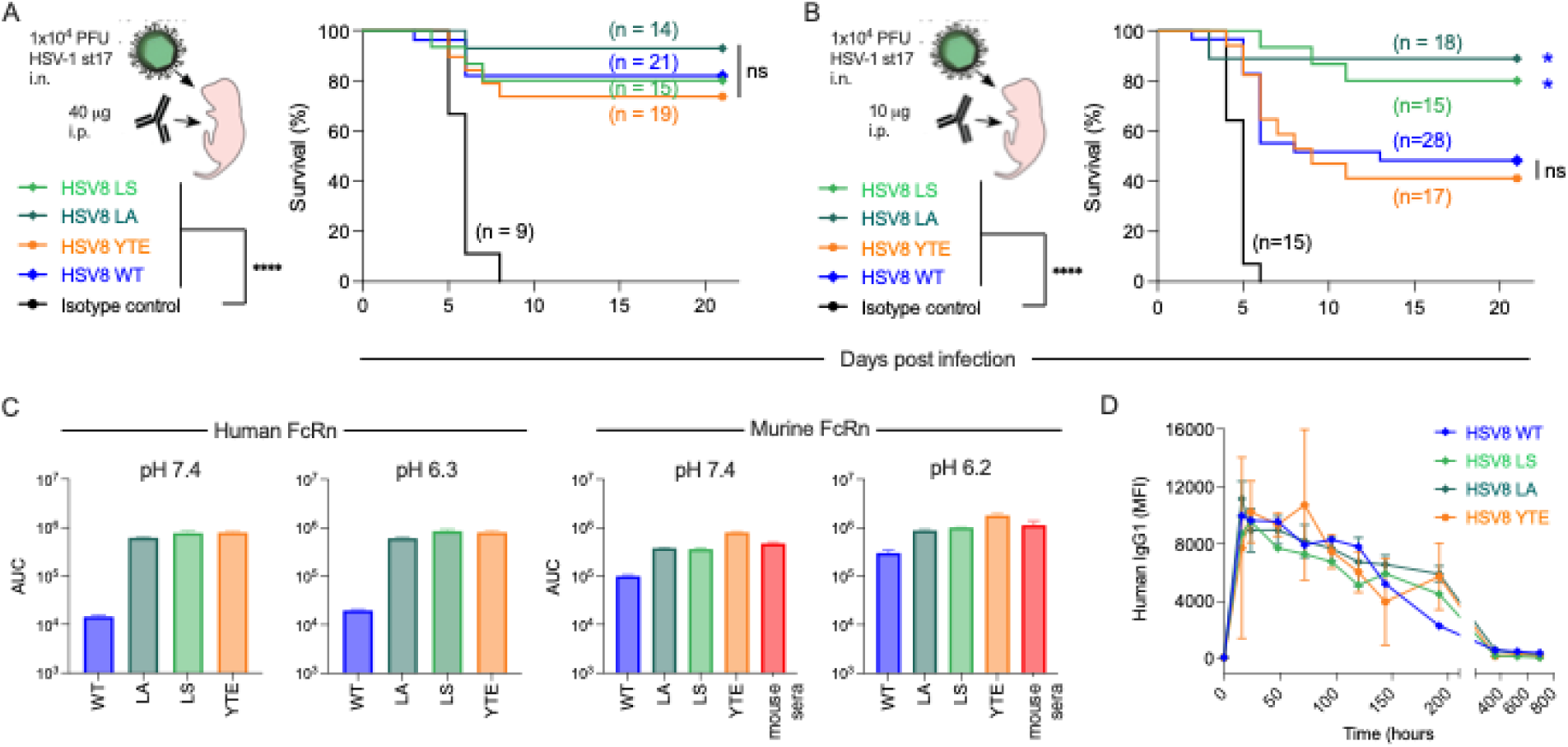
Improved protection against lethal HSV-1 challenge is afforded by human half-life engineered HSV8 mAb without improved pharmacokinetics. **A-B**. Survival of pups following mAb treatment. Immediately before lethal intranasal (i.n.) challenge with 1×10^4^ plaque forming units (PFU) of HSV-1 st17, 2-day-old C57BL/6J mice were given 40 µg (**A**) or 10 µg (**B**) mAb by intraperitoneal (i.p.) injection. The number of mice in each condition is reported in inset. Statistical significance as determined by the log-rank (Mantel-Cox) test (^∗^p < 0.05, ^∗∗∗∗^p < 0.0001) for each mAb compared to isotype control is reported in legend and between HSV8 WT and variants in inset. **C**. Area under the curve (AUC) of binding of HSV8 WT and its variants to human (left) and mouse (right) FcRn. Error bars represent standard deviation from the mean of technical replicates. **D**. Pharmacokinetic profile of HSV8 WT and variants following 10 µg i.p. mAb administration in adult C57BL/6J mice (n=3 per mAb).

We therefore measured the ability of these mAbs to bind to human and mouse FcRn at both extracellular and near-endosomal pH (**Figure 1C, Figure S2A**). As expected, HSV8 YTE, LA, and LS all displayed improved binding to human FcRn at low pH. As has been reported previously *(48)*, HSV8 YTE showed somewhat greater binding to mouse FcRn than HSV8 WT, but overall, each of the engineered mAbs exhibited similar binding to IgG derived from mouse sera. To directly evaluate the potential of differences in mAb bioavailability to influence survival, we evaluated mAb PK in serum following i.p. administration of a 10 µg dose in adult C57BL/6J mice. There was no difference in mAb kinetics between any of the four Fc variants (**Figure 1D**), as previously reported (*48, 53, 54)*. Taken together, these data indicate that the survival benefit observed for HSV8 LS and HSV8 LA is independent of improved half-life in mice and an additional property must explain these findings.

### Functional Characterization of HSV8 half-life variants

As these half-life mutations may also impact FcγR affinity and effector functions, we characterized the ability of these antibodies to bind to human and mouse Fcγ receptors. HSV8 WT, LA, and LS all bound equivalently to the activating Fcγ receptors: human FcγRIIA, human FcγRIIA, murine FcγRIII and murine FcγRIV; HSV8 YTE, in contrast and as expected, displayed reduced binding to all Fcγ receptors tested **(Figure S2B**). Consistent with receptor binding profiles, all variants of HSV8 were able to mediate effector functions such as ADCC, phagocytosis, and complement deposition in simplified assays, and some of these activities were somewhat reduced for HSV8 YTE (**Figure S2C)**.

In contrast to their comparable Fcγ receptor binding and effector function profiles, HSV8 LA, LS, and YTE demonstrated improved neutralization potencies against multiple HSV-1 and HSV-2 strains as compared to WT IgG1 (**Figure 2A**). HSV8 as a WT IgG1 molecule is a potently neutralizing antibody, with a 50% effective concentration (EC_50_) of 1.01 nM against HSV-1 st17 as measured by plaque reduction. However, LS, LA, or YTE mutations in the Fc region, typically considered to be independent of antigen binding, improved neutralization and reduced the EC_50_ of these mAbs, in some cases to sub-nanomolar ranges (**Figure 2B**). Variation in the degree of enhancement afforded Fc variants was observed across strains, with a particularly substantial impact on the neutralization of HSV-2. This result was unexpected, as modifying the Fc region is not canonically thought to impact Fab-dependent functions such as viral neutralization. Indeed, testing of mAb binding to recombinant soluble and cell surface-expressed gD was comparable across variants (**Figure 2C**). However, differential binding of both HSV8 WT and LA forms as well as isotype control mAb in WT and LS form was observed for virally-infected cells (**Figure 2D**). As a first indication of the potential of vFcR-mediated ABB to result in decreased susceptibility to mAb, the half-life extension mutation LA resulted in increased levels of infected cell surface binding of HSV8 (**Figure 2E**). In contrast, the low level of binding observed for the isotype control was reduced by the half-life extension mutation LS.

**Figure 2:**
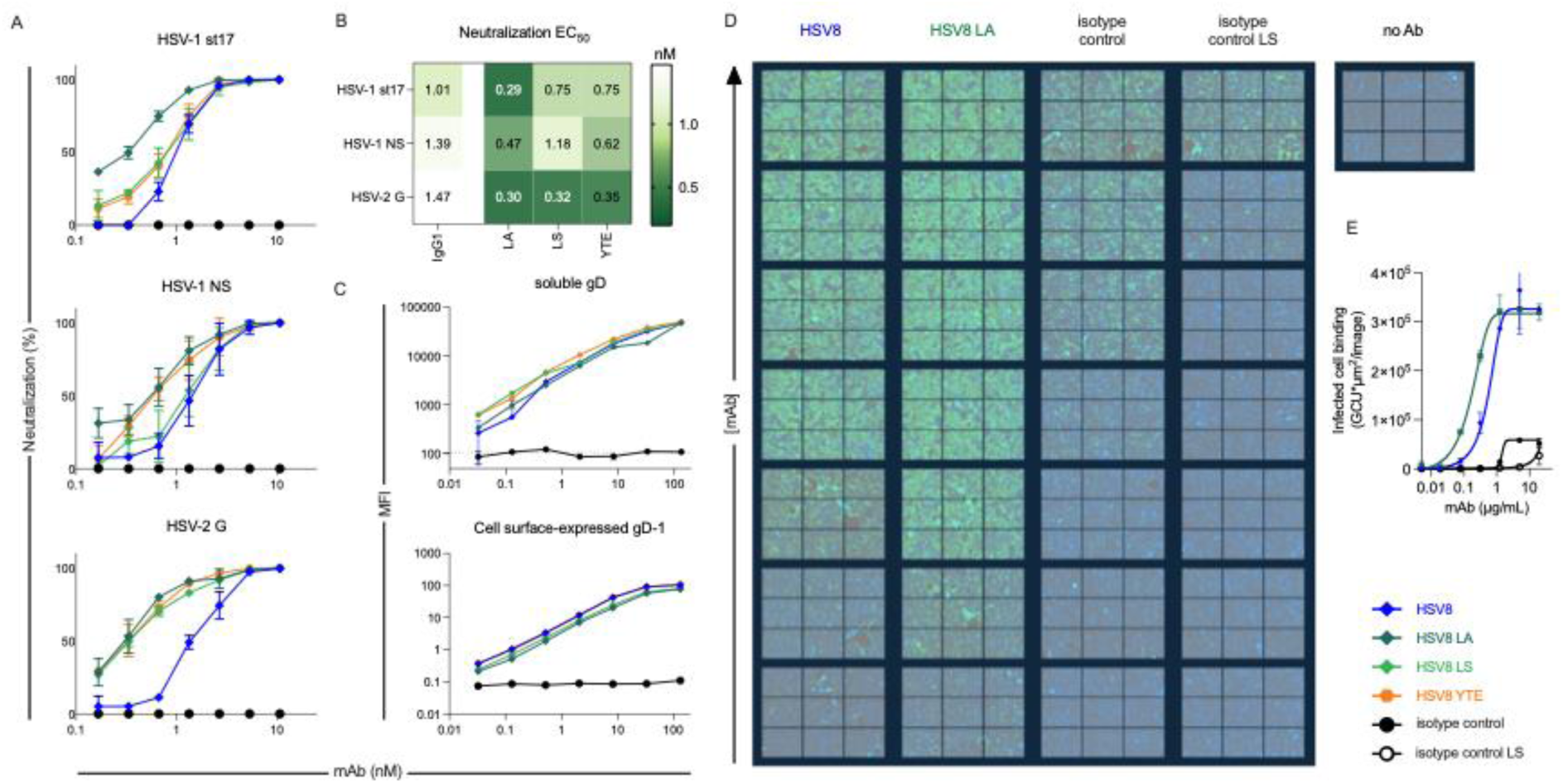
FcRn affinity mutations impact mAb neutralization and infected cell binding. **A.** Ability for HSV8 variants to neutralize HSV-1 st17 (top), HSV-1 NS (center), and HSV-2 G (bottom) by plaque reduction assay. **B**. Heatmap of neutralization midpoint Effective Concentration (EC_50_) values for each HSV8 variant and virus strain. **C**. Ability for HSV8 WT and variants to bind to recombinant gD conjugated to magnetic beads (top) or cell-surface expressed HSV-1 gD (bottom). **D**. Immunofluorescence images acquired at 20x magnification of IgG (green) bound to the surface of HSV-1 st17-infected cells (nuclei stained blue) across a titration range. **E**. Quantitation of infected cell binding. Error bars represent standard deviation from the mean. MFI – median fluorescent intensity. Assays were performed in technical and biological replicate.

### Human half-life variants impact HSV-1 gE binding

We next wanted to more directly address the hypothesis that these half-life extension mutations on the Fc region may impact binding to gE, as FcRn and gE recognize overlapping epitopes. This hypothesis is consistent with the crystal structure of the gE:Fc complex*(37)* **(Figure 3A**). The YTE set of mutations at positions 252, 254, and 256 in the CH2 domain of the Fc region all map directly to known interaction site with gE. Additionally, LS and LA mutations at positions 428 and 434 in the CH3 domain also flank one of the most important gE contact points His435 *(40)*. To test this hypothesis, we recombinantly produced soluble gE from HSV-1 st17 and measured gE binding to WT and engineered mAbs. We first measured the ability for gE to bind antibody-antigen complexes, mimicking ABB, in which Ab is simultaneously bound to gD through the Fab domain, and gE through Fc. HSV-1 gE bound WT HSV8 but was unable to bind HSV8 LA, LS, or YTE (**Figure 3B**). Like FcRn, HSV-1 gE is reported to exhibit pH-dependent binding *(55)*. As expected, at low pH, we observed reduced binding to the WT IgG1 mAb, while the inability of HSV8 LA, LS, and YTE to bind was unchanged (**Figure 3C**).

**Figure 3:**
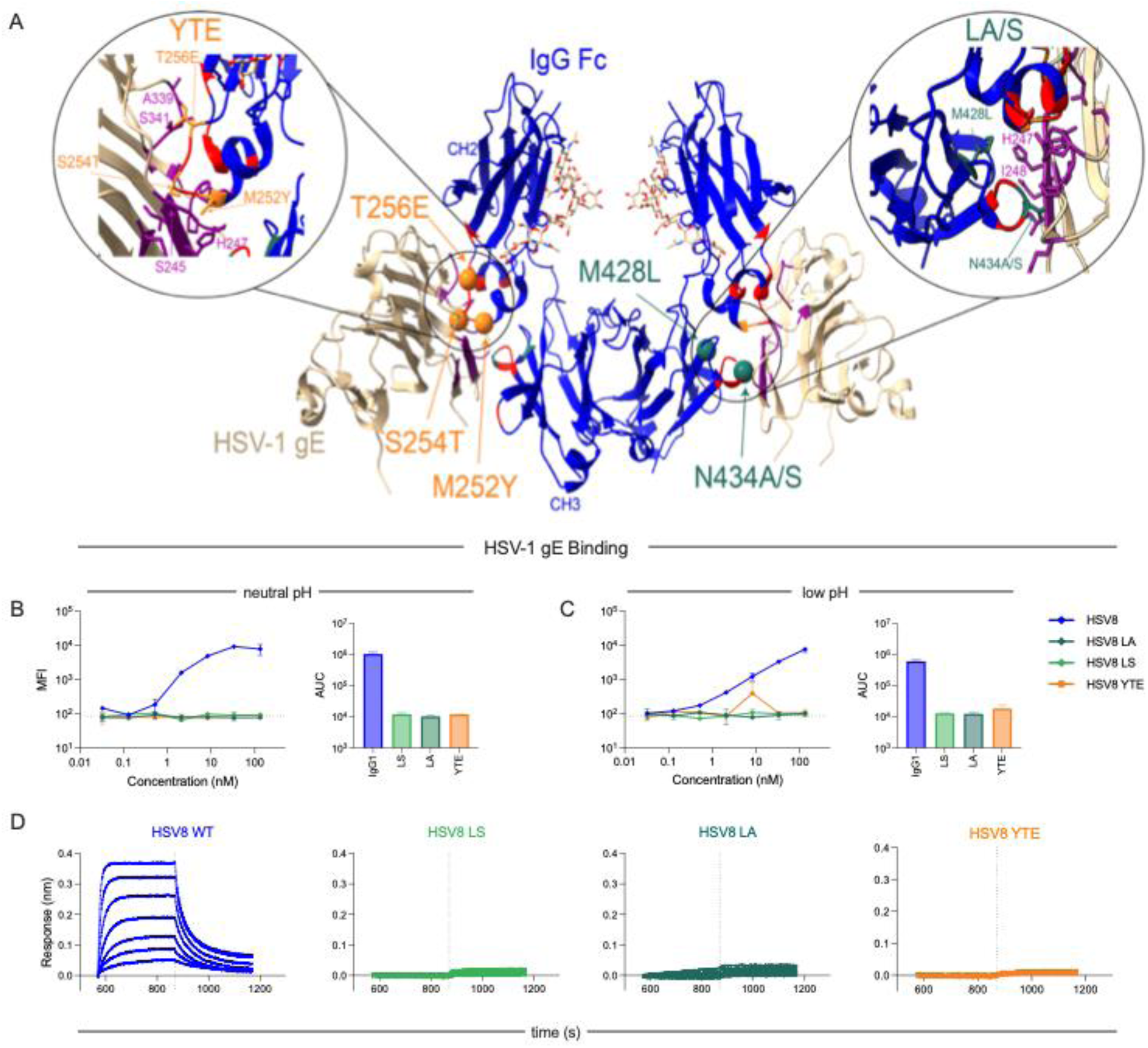
Half-life extension mutations impact HSV-1 gE binding. **A.** Visualization of half-life mutations modeled on the co-crystal structure of the HSV-1 gE (tan) and IgG Fc (blue) complex (pdb: 2jg7). gE contact residues on IgG Fc are colored in red, Fc contact residues on gE are in purple. YTE mutations are colored in orange and LS/LA mutations in green. Insets detail the contact regions and mutated residues. **B-C**. Ability for tetramerized HSV-1 gE to bind to the Fc region of gD-bound mAb at pH 7.2 (**B**) and 6.15 (**C**). Area under curve (AUC) values are plotted in inset. **D**. Association and dissociation curves of HSV8 variants over a concentration range binding to HSV-1 gE as measured by Biolayer Interferometry. Assays were performed in technical and biological replicate.

Lastly, for a more sensitive and label-free readout of the protein-protein interactions between HSV-1 gE and each IgG1 mutant, we turned to biolayer interferometry (BLI) to define the kinetic profile of association and dissociation (**Figure 3D**). WT HSV8 showed characteristic binding curve of concentration-dependent association with gE with a fast on- and fast off-rate profile. In contrast, no binding was observed for HSV8 LA, LS, or YTE. We hypothesized that elimination of gE binding disrupts this mechanism of immune evasion, resulting in improved viral neutralization and *in vivo* protection.

### HSV-1 gE discriminates between IgG3 allotypes

Natural human IgG subclasses and allotypes also differ in binding to human FcRn and prior reports demonstrate that the vFcR displays both species and human IgG subclass specificity (*38, 56)*. We therefore measured gE binding to several allotypes of human IgG3 *(57)* as well as murine IgG2a via BLI (**Figure 4A**). Consistent with prior work *(56)*, HSV-1 gE did not bind murine IgG2a, but allotypes of human IgG3 demonstrated differential binding to HSV-1 gE. Of the six allotypes selected for testing on the basis of sequence variation in their predicted contact residues, four displayed no binding to gE while two retained binding. This difference could be explained by amino acid sequence differences at two known gE interaction residues, His435 and Tyr436 *(37)* on IgG Fc (**Figure 4B**). IgG3m15* and IgG3m16*, the two allotypes that bound HSV-1 gE, differ from the other IgG3 allotypes but match human IgG1 at these positions (**Figure 4B**). Allotypes of human IgG3 with H435R replacement with or without Y436F mutation did not bind HSV-1 gE, pointing to their differential importance. Overall, these data align with reported differences among IgG3 allotypes for HSV-1 vFcR inferred from experiments evaluating binding to infected cells *(58)*, but extend prior work by providing direct kinetic analysis of protein-protein interactions. This allotype-dependent profile is shared with *Staphylococcal* Protein A binding to IgG Fc *(58)*, potentially pointing to broader benefits of this variation that, all else being equal, is expected to result in reduced steady state serum IgG3 levels.

**Figure 4:**
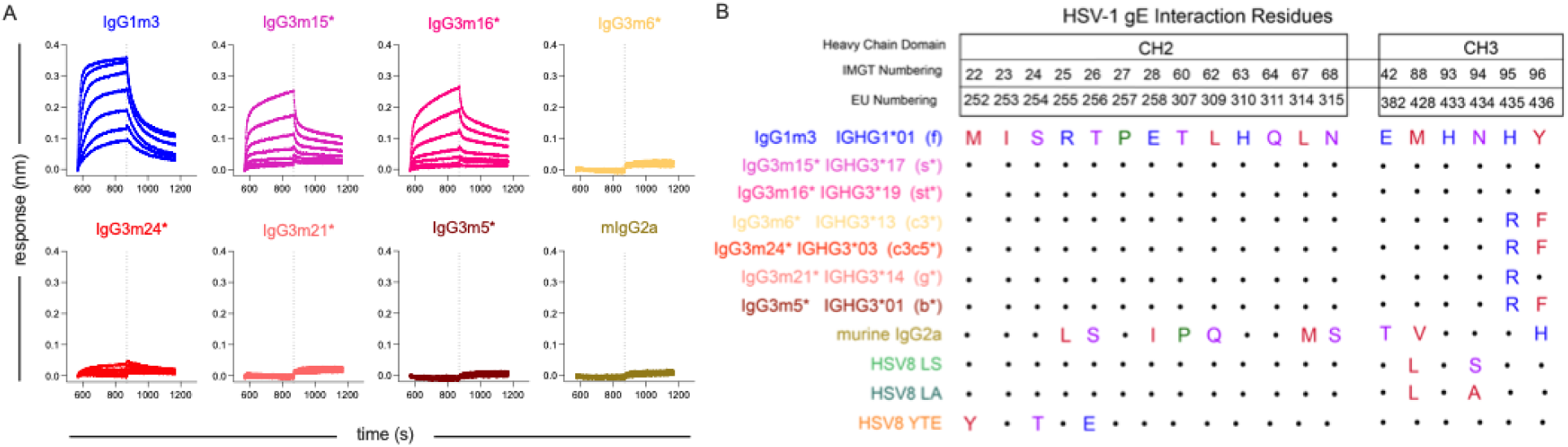
HSV-1 gE discriminates between species and human IgG3 Allotypes. **A.** Association and dissociation curves of human IgG1, human IgG3 allotypes, and murine IgG2a binding to HSV-1 gE as measured by Biolayer Interferometry. **B**. IgG Fc domain sequence alignment of known gE contact residues for each mAb and IgG allotype tested. Dots represent consensus as compared to human IgG1 (IgG1m3). Amino acid residues are colored by their chemical properties.

### Natural HSV-1 vFcR-evading mAbs show improved neutralization activity

Given the similar patterns in HSV-1 gE binding associated with natural IgG variation in human populations and engineered versions of HSV8, we tested non-binding murine IgG2a, human IgG3m15* (s*, binding), and IgG3m5* (b*, non-binding) in neutralization assays to define the importance of natural variation in vFcR activity to the antiviral activity of antibodies induced by vaccination or natural infection. HSV8 variants lacking vFcR binding displayed greater neutralization potency for HSV-1 NS as compared to HSV8 WT (**Figure 5A**). Among natural human IgG variants tested, IgG3 b* was most potent, and IgG3 s* the least.

**Figure 5:**
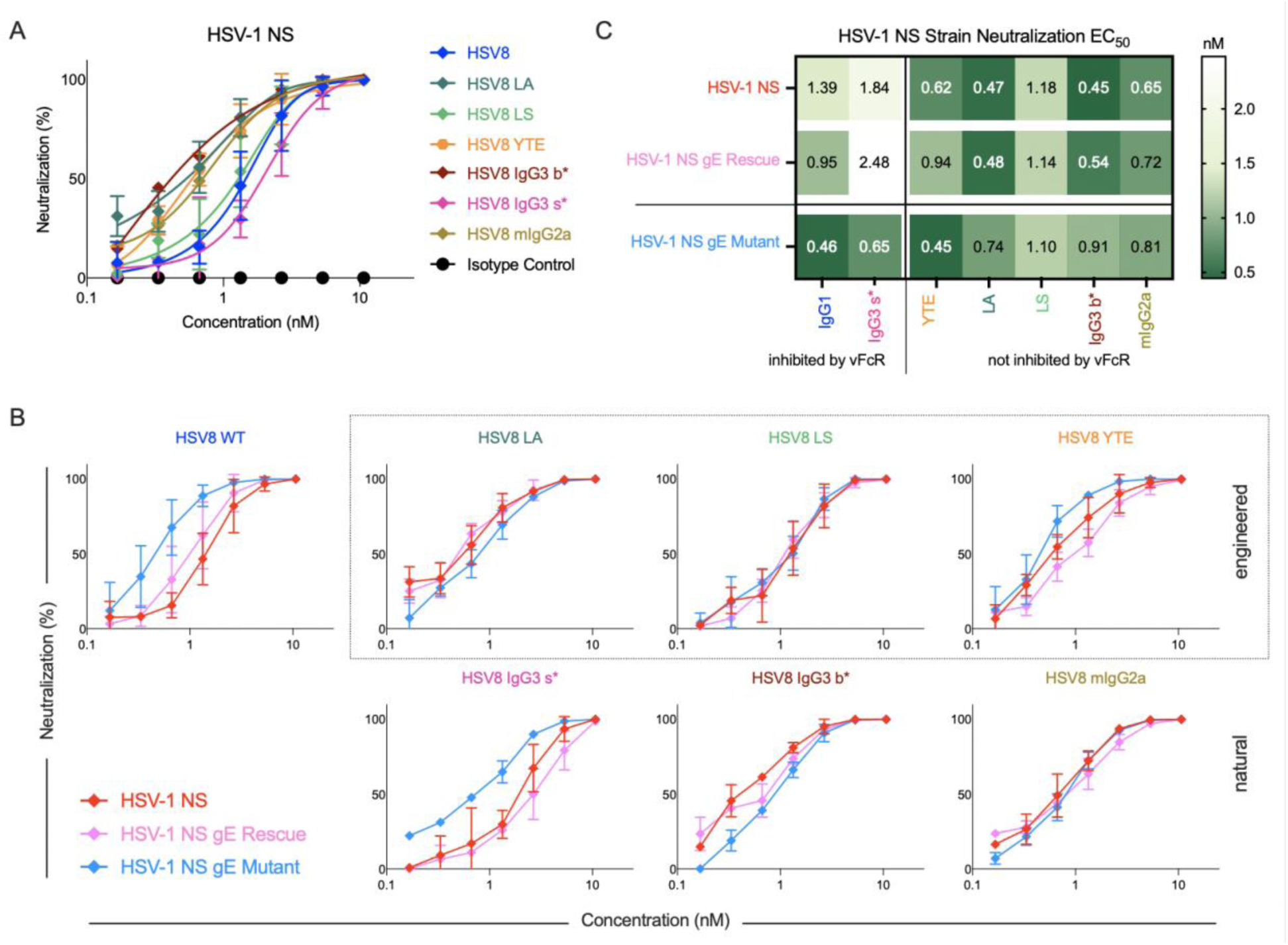
Eliminating gE-Fc interaction improves mAb neutralization for engineered and natural vFcR evading antibodies. **A.** Neutralization curves for engineered, species-, and subclass-switched HSV8 mAbs tested against WT HSV-1 NS by plaque reduction assay. **B**. The ability for HSV8 mAb panel to neutralize HSV-1 NS (red), as compared to a non-IgG binding HSV-1 NS gE mutant (blue), and reverted HSV-1 NS gE rescue strain (pink) as measured by plaque reduction assay. **C.** Heatmap of neutralization EC_50_ values observed for each HSV8 variant and virus tested. Dots represent the mean and error bars represent standard deviation. Neutralization assays were performed in technical and biological replicate.

### gE mutant virus is insensitive to vFcR-evading variants of HSV8

As a complement to mAb modifications, we next turned to viral genetics to address the impact of vFcR activity by using the HSV-1 NS gE264 (gE mutant) that contains a four amino acid insertion in gE that disrupts IgG Fc binding. This mutant strain, however, largely maintains other gE activities and viral pathogenesis *(39)*, unlike gE null strains that are largely non-pathogenic in mice (*30, 59, 60)*. In contrast to the vFcR-binding mAbs HSV8 WT IgG1 and IgG3 s*, which exhibited potent neutralization of the gE mutant virus, HSV8 LA, LS, YTE, mIgG2a, and IgG3 b* neutralized the parental NS, gE mutant, and gE revertant strains similarly well both qualitatively (**Figure 5B**) and quantitatively (**Figure 5C**). Overall, viruses with intact gE were resistant to neutralization by HSV8 variants that can undergo ABB (IgG1 and IgG3 s*). In contrast, neutralization activity of HSV8 variants that lacked gE binding were unaffected by gE vFcR activity.

### Eliminating vFcR:Fc interactions improves mAb-mediated protection in neonatal mice

We next defined the relevance of viral vFcR activity to HSV8 antiviral efficacy across natural and engineered Fc variants *in vivo* using the vFcR-modulated HSV-1 NS strains. For these strains, HSV8 WT at a dose of 10 µg was sufficient to provide a high level of protection (**Figure S3**). However, at 5 µg, both HSV8 LA and HSV8 LS provided substantially (∼80 and 40% survival, respectively) and significantly better protection than WT antibody (∼5% survival) against a lethal HSV-1 NS challenge (**Figure 6A**). Despite evading gE binding, 5 µg HSV8 YTE provided similarly poor efficacy, potentially due to compromised effector function, and weights among surviving pups were similar (**Figure S4**).

**Figure 6:**
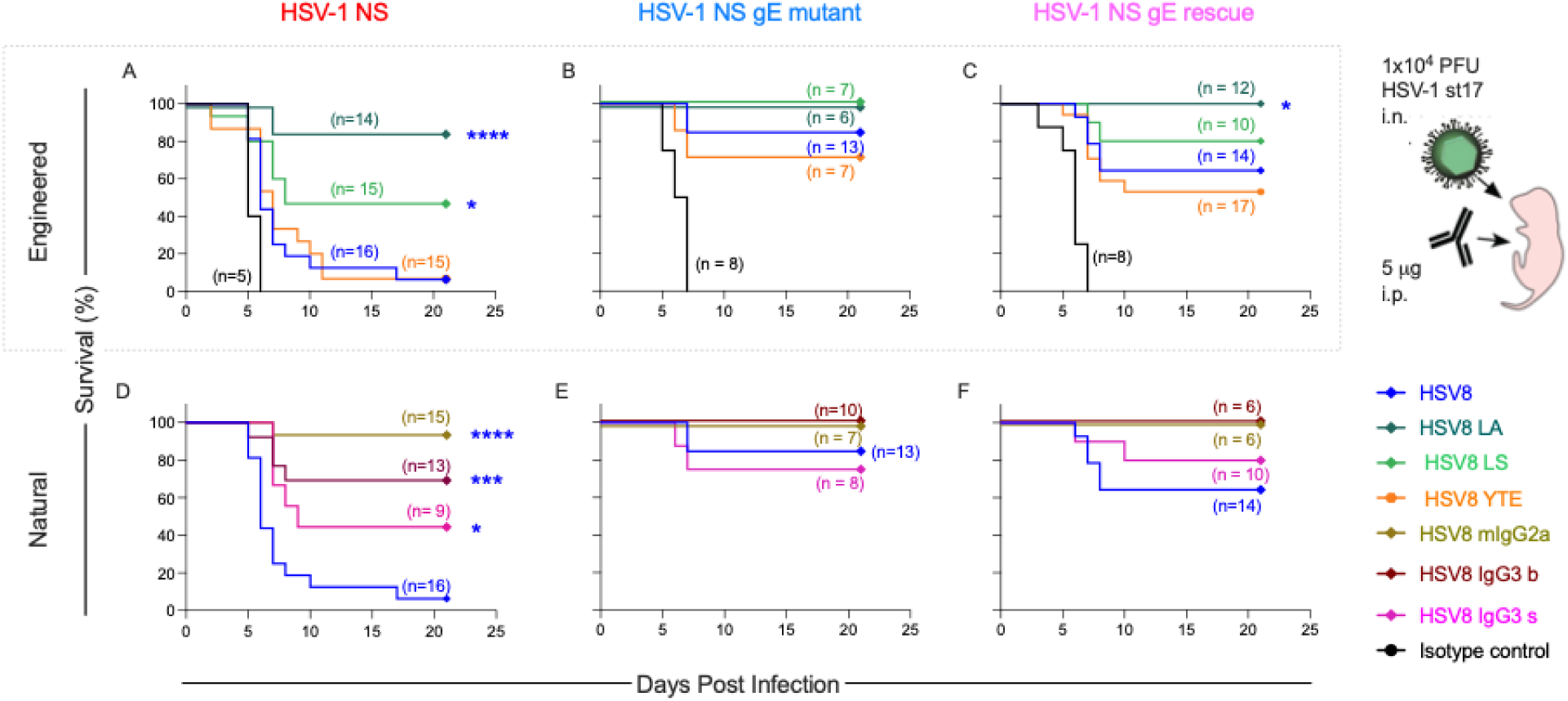
Natural or engineered evasion of HSV-1 gE:Fc interactions results in improved mAb potency *in vivo*. **A-F.** mAbs (5 μg) were delivered intraperitonially to two-day old pups immediately before a lethal (1×10^4^ PFU) challenge with HSV-1 NS (**A, D**), HSV-1 NS gE mutant (**B, E**), or HSV-1 NS gE rescue (**C, F**). Number of mice in each condition and statistical significance of indicated variant to WT HSV8 determined by the log-rank (Mantel-Cox) test (^∗^p < 0.05, ***p < 0.001, ^∗∗∗∗^p < 0.0001) are reported in inset.

When pups were instead challenged with HSV-1 NS gE mutant virus, treatment with any of these HSV8 variants resulted in >75% survival (**Figure 6B**), demonstrating the impact of vFcR activity on mAb efficacy. In contrast, pups treated with an isotype control mAb all succumbed to viral infection, demonstrating the maintained pathogenicity of gE mutant virus. While the HSV-1 NS gE rescue virus exhibited an intermediate rather than fully reverted phenotype, the same rank order of mAb activity to the WT NS strain was observed (**Figure 6C**). Overall, experiments with engineered virus and engineered HSV8 demonstrate that intact vFcR activity reduces mAb efficacy, and mAbs engineered to lack vFcR binding provide robust protection even at a low dose.

Natural vFcR-evading HSV8 variants (mIgG2a and IgG3b*) afforded a similar degree of protection as the LA and LS engineered forms across these virus strains (**Figure 6D-F**). Interestingly, the IgG3 versions of HSV8, regardless of susceptibility to the vFcR, provided significantly improved survival as compared to the WT IgG1 for the parental HSV-1 NS challenge (**Figure 6D**), suggesting more broadly enhanced antiviral activity of this subclass. In sum these results demonstrate that eliminating the ability for the vFcR to interact with the Fc region of an HSV-specific mAb results in improved neutralization potency and improved *in vivo* survival. Moreover, commonly used human half-life extension mutations that eliminate vFcR binding may provide a therapeutic advantage through both increasing half-life in humans and eliminating susceptibility to a key immune evasion mechanism by the virus.

### Kinetics of HSV gB and gC-specific IgG subclasses in humans

As subclass switching to IgG3 improved HSV8-mediated protection *in vivo* and the vFcR binds IgG1 but not IgG3 (*38, 61)*, we wanted to understand the kinetics of the IgG response to primary HSV infection in humans. We profiled the antibody response against three HSV surface glycoproteins, gD, gC, and gB, in a cohort of individuals with laboratory-documented primary genital HSV-1 infection at serial time points for 12 months postinfection. For all antigens profiled, we observed stereotypic increases in IgG1 responses over time (**Figure 7A-C, Figure S5**). Interestingly, unlike gB and gD, detectable IgG1 responses against gC were not observed at the first time point (2 weeks post study enrollment) (**Figure 7B, Figure S5B**), suggesting differential kinetics across glycoproteins. Unexpectedly, both gB and gC IgG3 responses slowly increased over time (**Figure 7B-C**), rather than arising early and waning over time as typical of acute viral infections *(62)*, and as was the case for gD. As the HSV vFcR cannot bind most allotypes of IgG3, these unconventional IgG subclass kinetics may be relevant for improved viral control against HSV.

**Figure 7:**
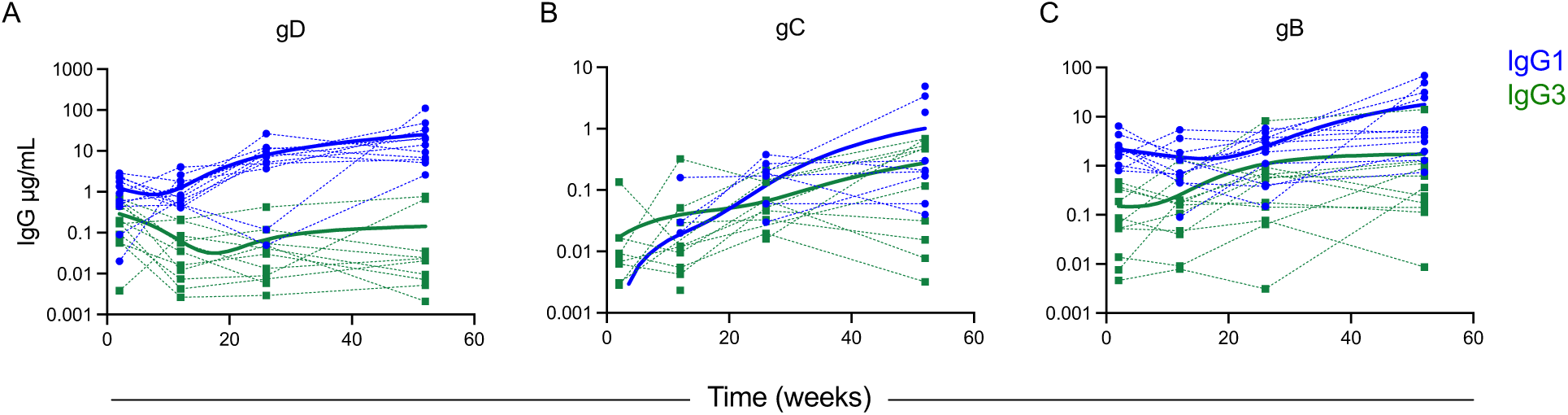
HSV-1 gB and gC display altered IgG subclass kinetics over time. **A-C.** Antibody responses to HSV-1 gD (**A**), gC (**B**), and gB (**C**) were measured in a cohort (n=11) of individuals with primary HSV-1 genital infection over the first year of infection. Population means for IgG1 (blue) and IgG3 (green) responses are shown with a thick line, and individuals with thin lines.

## Discussion

In the last decade, mAbs have rapidly emerged as important biologics for the treatment and prevention of infectious disease. Specifically, mAbs have been successfully used to treat acute viral infections such as Ebola, SARS-CoV-2, and RSV, the latter in neonates *(63–67)*. Additionally, mAbs with half-life extension mutations have not only been used, but also are safe in infants *(68–70)*. The clinical value of mAbs for other indications in neonates suggests that mAb treatment for nHSV should be explored to improve outcomes for nHSV infection. This study highlights an avenue for engineered mAbs to treat or prevent otherwise devastating nHSV infections. Moreover, it also establishes the potential importance of IgG3 subclass in mediating protection from HSV infection, in part by avoiding viral immune evasion mediated by gE, an attribute that may be important for vaccine development.

However, the use of mAbs in humans for either prophylaxis or therapeutic purposes for chronic viral infections caused by the Herpesviridae is a relatively nascent area of study. These viruses are extremely successful pathogens due to their life cycle and mechanisms of immune evasion — posing notorious challenges to vaccine development. Notably, many chronic viral infections have evolved Fc binding proteins to evade humoral immunity, broadly indicating that evasion of the Ab response is key to pathogen fitness. HSV-1, HSV-2, varicella zoster virus, and human cytomegalovirus (HCMV) each express Fc binding proteins that aid in immune evasion *(71)* and that may reduce the efficacy of mAb therapy. Fc engineering offers the possibility to evade vFcR while maintaining other mAb functions, though viral Fc binding proteins can share footprints with host FcRs and C1q (*36, 37)*, and mutating Fc residues can result in decreased effector function. This challenge was evident here, as HSV8 YTE eliminated binding of the vFcR, resulting in improved neutralization, but also reduced effector functions, resulting in a lack of improved *in vivo* efficacy relative to WT HSV8 mAb.

Recent studies have increased appreciation for the importance and role of Fc effector functions in mediating protection against herpesvirus infections(*8, 72–74)*. Our understanding of how natural IgG subclasses and allotypes differ in their activity against HSV not only provides insight into the Ab response to natural infection, but also informs vaccine design. IgG3 is functionally distinct from IgG1, and can better promote effector function and neutralization of some viruses *(75–77)*. Few studies have directly compared the efficacy of IgG3 and IgG1, but in HIV, IgG3 mAbs have demonstrated greater neutralization potencies *(78)*, improved Fc-mediated functions *(79)*, and phagocytic activity (*75, 80)*, each of which can contribute to viral clearance. Relative to IgG1, IgG3 has also demonstrated greater complement activity and bacteriolysis of *Neisseria meningitidis (81)*, and IgG3 versions of an IgG1 antibody against *Streptococcus pneumoniae* provided greater *in vivo* protection in mice *(82)*. Together, the expected improvement in effector functions of IgG3 combined with a natural lack of vFcR binding makes this subclass an attractive option. Indeed, we observed that IgG3 versions of HSV8, regardless of vFcR binding, provided improved efficacy as compared to IgG1.

Canonically, IgG3 responses against viral antigens arise early in infection and wane, sometimes rapidly, over time (*62, 83, 84)*. This kinetic profile is consistent with the arrangement of the human IgH locus and the radical biology of class-switching, in which upstream subclasses are excised from the genome. In contrast to these previous observations, we observed low or slowly increasing IgG3 responses to HSV-1 gB and gC over the course of primary genital HSV-1 infection. While gD was not among these antigens, the improved efficacy of HSV8 that resulted from subclass-switching to IgG3 provides evidence that HSV may have evolved to alter the typical induction of this “early responder” subclass. Consistent with our data for HSV, IgG3 responses in humans to HCMV rise over time for some antigens (gB and pentamer) but not others (tegument) *(85)*. Like HSV, HCMV encodes Fc binding proteins *(86)* that serve to evade humoral immunity (*87, 88)*. While HSV encodes a single vFcR, HCMV encodes four – gp34, gp68, gpRL13, and gp95. Of these, gpRL13 and gp95 cannot bind to the Fc region of IgG3 *(86)*, however the precise affinities, binding sites on IgG Fc, and roles in viral pathogenesis for all four vFcRs have yet to be described. Despite this knowledge gap, the fact that gpRL13 and gp95 cannot bind IgG3 is interesting given the atypical longitudinal profile of IgG3 directed against HCMV gB and pentamer throughout infection.

Further study will be needed to investigate IgG profiles in a larger number of individuals and in response to primary HSV-2 infection, nHSV infection, and in association with host IgG allotypic diversity. Similarly, while the efficacy improvements afforded by Fc engineering and subclass-switching are exciting and improve our understanding of antibody-mediated protection against HSV infection, we evaluated only one mAb, HSV8, which targets gD. Other mAbs, including those targeting gB have also shown efficacy in preclinical models and are being evaluated clinically (*10, 12)*. If Fc modification is a general means to improve *in vivo* efficacy, the clinical prospects of these mAbs might likewise improve. Understanding the mechanistic basis for how Fc modifications affect mAb efficacy will provide a basis for engineering more effective antibodies. Lastly, while we observed greater neutralization potencies of vFcR-evading HSV8 variants against HSV-2, we did not evaluate efficacy *in vivo* for HSV-2. To the best of our knowledge, no equivalent to the HSV-1 gE mutant virus used here exists for HSV-2, posing a barrier to dissection of the impact of vFcR activity in mAb-mediated protection against HSV-2.

In sum, this study demonstrates that HSV vFcR activity may be an important consideration in evaluating mAb therapeutics and antibody responses to infection or vaccination. vFcR activity, mediated by gE:Fc interactions, directly influenced both mAb neutralization potency and *in vivo* efficacy, reducing the ability for HSV8 IgG1 to protect neonatal mice from HSV-mediated mortality. Both natural and engineered Fc variants of HSV8 that lacked vFcR binding were more potently neutralizing and displayed improved efficacy. Collectively, these results demonstrate that the uncoupling of the HSV vFcR binding to IgG Fc is possible, leading to a generalized approach for creating more potent therapeutic antibodies.

## Materials and Methods

### Experimental Design

The rationale for this study was based on previous studies (*7, 8)*. Overall, we wished to determine the impact of the HSV vFcR on mAb-mediated protection and whether Fc engineering to eliminate vFcR interactions produced a more potent therapeutic. Sample sizes were determined from previous work using HSV-specific mAbs in the mouse model of nHSV infection. One to four litters are represented in each graph with the exact n listed in the inset for each survival curve. Endpoint criteria for the survival experiments were defined as excessive morbidity (hunching, spasms, and/or paralysis) and/or >10% weight loss from the previous measurement. Animal procedures were performed in accordance with Dartmouth’s Center for Comparative Medicine and Research policies and following approval by Institutional Animal Care and Use Committee (Protocol number: 00002151 – approved 240809).

### Human Samples

All participants provided written informed consent, and the study was approved by the University of Washington Human Subjects Division. Participants with virologically confirmed first episode genital HSV-1 infection were followed for one year as previously described *(89)*. A subset of participants (n=11) with negative HSV Western Blot at the screening visit, confirming primary HSV-1 infection, were enrolled into an immunologic substudy, with serial blood draws at defined time points (2 4, 6, 12, 26 and 52 weeks).

### Mouse procedures and viral challenges

Wild Type C57BL/6J (B6) mice were either purchased from The Jackson Laboratory or bred in house in accordance with Institutional Animal Care and Use Committee protocols. mAbs were administered via the intraperitoneal route with a 25µL Hamilton Syringe in a 20µL volume under 1% isoflurane anesthesia. The wild-type viral strains used in this study were HSV-1 st17syn+*(90)*, HSV-2 strain G*(91)* (provided by Dr. David Knipe), and HSV-1 NS (provided by Dr. Harvey Friedman)*(92)*. HSV-1 NS gE264 (HSV-1 gE mutant) and its revertant (HSV-1 gE rescue) were also provided by Dr. Harvey Friedman*(39)*. Viral stocks were prepared using Vero cells as described previously(*93, 94)*. Newborn pups were infected intranasally (i.n.) on day 2 postpartum with 1×10^4 PFU of the indicated HSV-1 strain in a 5µL volume under isoflurane anesthesia. Pups were then monitored for survival and infected pups were weighed daily through day 21 post infection. Endpoint criteria for the survival experiments were defined as previously described *(8)*.

### Viral Neutralization

Viral neutralization of all HSV strains was performed via plaque reduction neutralization tests *(8)*. Briefly, serially diluted mAb and 50 PFU of HSV-1 st17, or HSV-2 G, or HSV-1 NS, or HSV-1 NS gE mutant, or HSV-1 gE rescue were incubated for 1 hour at 37°C before being added to confluent vero cells grown in 6 well plates (Corning). Plates were incubated for 48 (HSV-1 st17) or 72 hours (HSV-2 G, HSV-1 NS, HSV-1 NS gE mutant, HSV-1 NS gE rescue). Viral neutralization (%) was calculated as [(number of plaques in virus only control well - number of plaques counted at mAb dilution)/# of plaques in virus only control well]*100. Assays were performed in technical and 2-3 biological replicates. EC50 values were calculated using a 4 point sigmoidal curve fit in Prism 10.

### Infected Cell Binding

A confluent monolayer of vero cells in a 96 well plate was infected with HSV-1 st17 at a multiplicity of infection of 1 and incubated for 18 hours at 37°C with 5% CO2. Media was removed and cells were washed with PBS before being fixed with 4% PFA. Fixed cells were then permeabilized with 0.1% Triton X-100 in PBS. Vero cells were washed and then blocked with 2.5% Normal Goat Serum (NGS) in PBS with 0.1% Triton X-100. Cells were washed and then incubated with serially diluted HSV-specific mAbs or isotype control for 1 hour at RT. Cells were washed once again before being incubated with a FITC-conjugated goat anti-human IgG (H+L) secondary antibody and 1µM TO-PRO-3 Iodide nucleic acid stain for 1 hour at room temperature. Cells were washed a final time before being imaged with an Incucyte (Sartorius) at 20x with 9 images per well. Green Intensity (GCU x µm^2/image) was calculated using Incucyte Software (Sartorius)

### Measurement of HSV-1 gE binding via Biolayer Interferometry (BLI)

50nM of biotinylated HSV-1 gE was immobilized onto streptavidin biosensors (Sartorius) in 1x Kinetics Buffer followed by association and dissociation measurements in 2-fold diluted mAb (15.63 nM to 1000nM) on an OctetRed96 instrument (ForteBio). Association and dissociation were measured for 300 seconds respectively. Binding curves were calculated following normalization to association baseline.

### Statistical Analysis

Prism 10 (GraphPad) was used for all statistical analyses. For survival studies, HSV-specific mAbs were compared to the isotype control and HSV8 Variants were compared to the WT IgG1 using the Log rank Mantel-Cox test. For human data, trendlines over time were generated using the LOWESS curve fit method for each IgG subclass for a given HSV-specific antigen.

## Acknowledgements

We would like to thank all members of the Ackerman and Leib labs for their support and invaluable feedback and scientific discussion. We acknowledge Andrew Hederman for assistance in developing the Biolayer Interferometry experiments and Audra Charron for assistance with immunofluorescence assays. We thank Zabb Bio for providing HSV8, HSV8 LA, HSV8 LS, and HSV8 YTE. We thank Harvey Friedman for providing HSV-1 NS, HSV-1 NS gE264, and HSV-1 rNS gE264.

## Funding

These studies were partially supported by National Institutes of Health grants, NEI 5R01EY009083-28 to D.A.L., NIAID grants 5P01AI098681-08 and 5R21AI147714-02 to D.A.L., U19AI145825 to M.E.A., R01AI176646 to D.A.L. and M.E.A., T32AI007363 to M.D.S, and R01AI178284 to A.M.S.

## Author Contributions

MDS, DAL, MEA conceptualized the study. MDS, IMB, NSK, CRG, USK, AMS conducted experiments. CJ provided human samples. DAL and MEA obtained funding and supervised the research. MDS generated figures and drafted the manuscript. MDS, DAL, MEA finalized the manuscript. All authors read through and edited the final version of the manuscript.

## Competing Interests

I.M.B., D.A.L., and M.E.A. report a patent, WO2020077119A1, for the use of HSV-specific mAbs as method for the treatment for nHSV infections. CJ reports consulting for GSK, Pfizer, and Assembly Biosciences and research funding from GSK and Moderna unrelated to this work. MEA reports consulting for Seromyx Systems and research funding from Moderna unrelated to this work.

## Data Availability

All data associated with this work are present in the main text and supplementary materials. Reasonable requests for reagents, resources, and data should be directed to the corresponding author and will be fulfilled with a materials transfer agreement.

