## Supplemental Materials for "Improving antibody-mediated protection against HSV infection by eliminating interactions with the viral Fc receptor gE/gI"

**Materials and Methods**

*HSV8 Pharmacokinetics*

Antibody pharmacokinetic experiment was performed as reported previously*(48)* with slight modifications. 10 µg of each HSV8 variant in a 100 µL volume was delivered to adult B6 mice of both sexes via the intraperitoneal route. Blood was taken via the retro-orbital sinus via a heparinized capillary tube starting at 16 hours post mAb administration and subsequently at 24 hours post administration and then approximately every 24 hours for the first 6 days. Blood was then drawn on a weekly basis either via the submandibular vein or via the retro-orbital sinus. Blood was spun at 2000xg for 20 minutes at 4 °C to separate plasma from other blood components. Plasma was separated and stored at -20˚C until quantification. Relative antibody amounts were quantified via a bead based assay *(76)*. Briefly, HSV-1 gD *(94)* and F(ab')2-goat anti-human IgG Fc (ThermoFisher A24478) were amine coupled to magnetic fluorescently barcoded microspheres *(95)*. Samples were diluted in assay buffer (0.1% bovine serum albumin (BSA) with 0.05% Tween-20 in PBS) prior to incubation with coupled microspheres. Bead-bound antibodies were detected with goat anti-human IgG PE (Southern Biotech) and detected via FlexMap 3D (Luminex) *(96)*.

*Monoclonal Antibodies*

HSV8 WT, LA, LS, and YTE mAbs were provided by ZabBio and Kentucky Bioprocessing. HSV8 was formatted with murine IgG2a and Kappa constant sequences using the VH and VL sequences as reported *(97)*. Recombinant DNA was synthesized by Twist Biosciences. HSV8 human IgG1 and Kappa Light Chain DNA sequences were also synthesized by Twist Biosciences and cloned into pCMV expression vectors. HSV8 IgG3 b* and s* allotypes were cloned by overlap extension PCR combining the HSV8 VH and IgG3 constant domains from plasmids containing the IgG3 allotypes *(57)*. Cloned expression plasmids containing HSV8 heavy, or light chains were sequence confirmed via Sanger sequencing. Other IgG3 allotypes used in this study have been reported previously *(57)*. Antibodies were expressed through co-transfection of the light chain and heavy chain plasmids in ExpiCHO cells (Thermo Fisher) according to the manufacturer's protocols. Ten days after transfection, cultures were spun at 8,000 x g for 90 minutes to pellet the cells and the supernatants were sterile filtered (0.22 µm). Murine IgG2a was purified using a custom packed Protein A (Cytiva) column and eluted with 100mM glycine pH 3.0 and then immediately neutralized with 1M Tris-HCl pH 8.0. Eluate was concentrated using Amicon 30kDa cut-off columns (Cat # UFC903024). HSV8 IgG3 allotypes were expressed in the same system but purified using Protein A/G Plus Agarose (Thermo Fisher). mAbs were passed over endotoxin removal columns (Thermo Fisher), aliquoted and snap-frozen before being stored at -80˚C for later use.

*Measurement of binding to human and mouse Fc receptors.*

Recombinant HSV-1 gD antigen *(94)* was provided by Dr. Gary Cohen (University of Pennsylvania) and coupled to magnetic microspheres *(95)*. HSV8 mAbs were serially diluted in assay buffer (0.1% BSA with 0.05% Tween-20 in PBS) and incubated overnight with antigen-coupled beads at 4 °C with constant shaking. Beads were then washed before being incubated with recombinant biotinylated Fc gamma receptors *(98)* (Duke Human Vaccine Institute), mouse Fc gamma receptors, or with human or mouse neonatal Fc receptors. For all experiments, Fc receptors were tetramerized with streptavidin-PE prior to incubation with antibody immune complexes. Fc receptors were incubated with antibody immune complexes for 1 hour at room temperature with constant shaking. For pH differences in FcRn binding, recombinant FcRn was tetramerized with streptavidin-PE in assay buffer at pH 6.0 and incubated with antibody immune complexes for 1 hour at room temperature with constant shaking. The beads were washed and subsequently analyzed via the FlexMap3D instrument (Luminex). The median fluorescent intensity of at least 10 beads/region was recorded. A buffer only control and an isotype control antibody were included to determine assay background and antigen-specificity, respectively. Area under the curve was calculated using Prism10 (GraphPad).

*Antigen Binding*

The ability for the HSV8 variants to bind to soluble recombinant HSV-2 gD was evaluated via Luminex *(8)*. Briefly, serially diluted HSV8 variants were incubated with HSV-2 gD magnetic microspheres overnight at 4 °C with constant shaking. Immune complexes were washed and incubated with goat anti-Human IgG PE (Southern Biotech) for 1 hour at room temperature with constant shaking. The beads were washed again and analyzed via the FlexMap3D system (Luminex). HSV8 variants were also evaluated for their ability to bind cell-surface expressed gD-1. HEK293Ts were engineered to express HSV-1 gD as a surface antigen *(8)*. gD-expressing 293Ts and non-transfected 239Ts (control) were trypsinized, washed twice with PBS, and 200,000 cells/well were added to a 96 well V bottom plate (USA Scientific). HSV8 variants were diluted to 20 μg/mL and serially diluted 4-fold in PBS with 1% BSA before being added to the cells. After a 1 hour incubation on ice, the cells were washed twice with 1% BSA in PBS and stained with 10 μg/mL AlexaFluor 647 goat anti-human IgG (H+L) cross-adsorbed secondary antibody (ThermoFisher). After a 30 minute incubation in the dark, cells were washed twice and then fixed with 4% paraformaldehyde. Antibody binding was quantified via a MACSQuant Analyzer (Miltenyi). The experiment had two biological replicates and the data was analyzed with FlowJo version 10.8.2.

*Measurement of HSV-1 gE binding to IgG Fc*

Recombinant gE ectodomain from HSV-1 st17 DNA was synthesized by Twist Biosciences and cloned into pCMV expression vectors. A 6x His and AviTag were included at the C-terminus of the ectodomain. The expression plasmid was sequence confirmed via Sanger sequencing. Soluble HSV-1 gE was expressed via transfection of Expi293 (ThermoFisher) cells in accordance with the manufacturer's protocols. 5 days post transfection, cells were harvested and spun at 4000xg for 1 hour and cell supernatants were sterile filtered (0.22 μm). Supernatant was passed over a custom-packed HisPur™ Ni-NTA (ThermoFisher) column. Bound gE antigen was eluted using 250mM imidazole in accordance with manufacturer's protocols. Eluted gE-1 antigen was concentrated, and buffer exchanged into PBS. Purity and size were confirmed via SDS-

PAGE gel and an ELISA using an HSV-1 gE specific mAb (Virusys Corporation). HSV-1 gE was biotinylated via BirA biotin ligase. The ability for gE to bind to IgG Fc was evaluated via a custom multiplex microsphere experiment like Fcγ receptor binding as described elsewhere in this paper. Briefly, HSV8 variants and other HSV gD-specific antibodies were incubated with gD-coupled magnetic microspheres overnight at 4 ˚C with constant shaking. Biotinylated HSV-1 gE was tetramerized with streptavidin-PE before being incubated with immune complexes for 1 hour at room temperature with constant shaking. Beads were washed and binding was analyzed via a FlexMap3D Instrument (Luminex). Buffer only wells and a mouse IgG2a antibody were used as negative controls. For pH dependent binding, gE was tetramerized and bound to IgG Fc at pH 6.15.

*Antibody Dependent Cellular Cytotoxicity*

A CD16 reporter assay was performed as previously described (100). Briefly, the wells of a high-binding 96 well plate (Corning) were coated with 1 µg/mL recombinant gD-2 protein in PBS and incubated overnight at 4˚C. Plates were then washed 3x with 0.5% Tween-20 in 1x PBS and then blocked with 2.5% BSA in PBS for 1 hour at room temperature. HSV8 variants were serially diluted in assay medium (RPMI with 10% FBS, 1% P/S and 1% non-essential amino acids) before being added to washed plates with 100,000 Jurkat Lucia NFAT CD16 cells/well (Invivogen). Antibodies and cells were incubated at 37˚C with 5% $CO_2$ for 24 hours. Plates were spun at 300xg for 5 minutes and then a 25 µL volume of the cell supernatant was removed and added to a new, opaque white 96 well plate. A volume of 75 µL of reconstituted QuantiLuc (Invivogen) substrate was added to the cell culture supernatant and luminescence was immediately read on a SpectraMax Paradigm plate reader (Molecular Devices) using a 1 second integration time. A kinetic read time of 0, 2.5 and 5 minutes was performed. Reported values are the averages of the three reads. Buffer only wells and an isotype control were used as negative controls. A cell stimulation cocktail containing 2 µg/mL ionomycin was used as a positive control. The assay was performed in technical replicate.

*Antibody-dependent cellular phagocytosis (ADCP)*

ADCP was performed as previously described *(101)* with slight modifications. Briefly goat anti-human IgG F(ab')2 was covalently coupled to yellow green 1µm carboxylate beads (ThermoFisher). Antibodies were diluted in culture medium (RPMI with 10% FBS and 55 µM beta-mercaptoethanol) and serially diluted 4-fold 7 times and incubated with anti-human IgG beads for 2 hours at 37˚C to form immune complexes. 25,000 THP-1 (ATCC) cells/well were added to the immune complexes and incubated at 37˚C for 4 hours. Cells were washed 2x with cold PBS prior to fixation with 4% paraformaldehyde. Cells were analyzed on a NovoCyte Advanteon flow cytometer (Agilent). A phagocytosis score was calculated as (percentage of FITC+ cells) x (the geometric mean fluorescence intensity (gMFI) of the FITC+ cells)/100,000. Buffer only wells were used as negative controls, and the assay was performed in technical replicate with two biological replicates.

*Antibody-dependent complement deposition (ADCD)*

HEK293Ts engineered to express HSV-1 gD on the cell surface were used as target cells to measure the ability of the HSV8 variants to mediate ADCD as reported previously *(8)*.

*Measurement of IgG1 and IgG3 against HSV-1 glycoproteins in human plasma via Multiplex binding antibody assay (MBAA)*

MBAA was performed as described elsewhere with some modifications *(102, 103)*. HSV-1 gB was provided by Drs. G. Cohen and R. Eisenberg (UPenn). gC1 and gD1 were expressed in 293E cells and purified using Ni-NTA agarose followed by size exclusion chromatography on Superdex 200 (Cytiva). HSV-1 proteins and control antigen, tetanus toxoid (Enzo, catalog number ALX-630-108) were conjugated to Bio-Plex Pro Magnetic COOH beads in a ratio of 10 µg of antigen per 2.5 x $10^6$ beads in a two-step carbodiimide reaction. First, beads were washed and resuspended in Activation Buffer (100 mM MES, pH 6) and then incubated with N-hydroxysulfosuccinimide (Sulfo-NHS, catalog number 24520; ThermoFisher) and 1-ethyl-3-[3-dimethlyaminopropyl]carbodiimide-HCl (EDC, catalog number 77149; ThermoFisher) also dissolved in Activation Buffer for 20 minutes on an end-over-end rotational mixer at room temperature protected from light. Activated beads were washed three times in Activation buffer. For coupling, antigen was mixed with activated beads and reaction was carried out for 2 h on a rotational mixer at room temperature protected from light. Conjugated beads were washed three times with Wash buffer (PBS, 0.05% Tween-20, 1% BSA, 0.1% $NaN_3$) and finally resuspended in Wash buffer at $10^7$ beads/ml. Beads were stored at 4 °C for no longer than 30 days.

Antigen-specific IgG was measured using two replicate dilutions. Beads were blocked with phosphate buffered saline (PBS; Gibco) containing 5% Blotto (Bio-Rad) and 0.05% Tween-20 (Sigma) and incubated for 1 hour with serially diluted plasma samples. Plasma pooled from ten HSV-2 seropositive and ten HSV-2 seronegative donors were used as positive and negative controls, respectively. After incubation with plasma samples, MagPlex beads were washed with PBST (PBS, 0.05% Tween-20) and incubated with either Mouse Anti-Human IgG1 Fc-PE (SouthernBiotech, catalog number 9054-09) or Mouse Anti-Human IgG3 Hinge-PE (Sothern Biotech, catalog number 9210-09). Finally, beads were washed 3 times and resuspended in PBS containing 1% BSA and 0.05% Tween-20. Median fluorescence intensity (MFI) by PE fluorescence for each bead type was collected on Bio-Plex 200 instrument (Bio-Rad) operated by Bio-Plex Manager v.6.0 software (Bio-Rad). The level of background was assigned by the MFI of antigen-conjugated beads incubated first with buffer (in place of serum), then with secondary antibody. Background MFI values for each antigen were subtracted from experimental measurements.

A standard curve run in duplicate was used to estimate IgG1 and IgG3 concentrations. For that, anti-human IgG Fab-specific (Southern Biotech) was conjugated to MagPlex beads. IgG-coupled beads were blocked, washed and incubated with serially diluted human standard human IgG (Sigma, catalog number I4506) for 1 h. Standard beads were washed and incubated with anti-human IgG-PE (SouthernBiotech, catalog number 2040-09), and MFI was measured as described above. MFI readings and associated IgG concentrations were fitted to a five-parameter logistic curve (5PL) using Bio-Plex Manager. A standard curve for each experiment was used to obtain the effective concentrations of IgG1 or IgG3 in plasma using the MFI measured with antigen-coated beads. Since plasma samples were also run as a dilution series, we used the mean of the estimated concentrations from the dilutions that yielded MFIs between 100 and 10,000. Plasma with all values above (below) this range were right (left) censored at the concentration of the minimum (maximum) MFI.

### Supplemental Figures and Legends

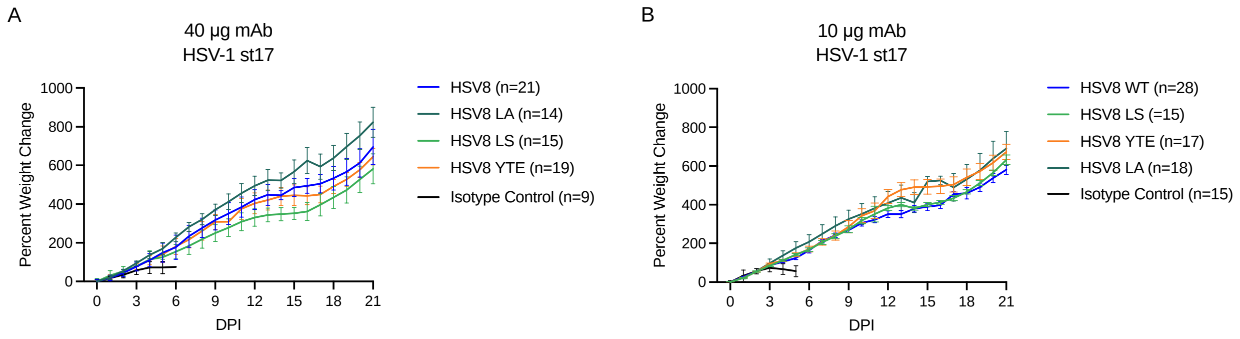


**Figure S1 (related to Figure 1). Percent weight change of mouse pups receiving HSV-specific mAbs. A-B**. Weight gain of pups following HSV-1 challenge. Immediately before lethal intranasal challenge with $1x10^4$ plaque forming units (PFU) of HSV-1 strain 17, mouse pups received 40 µg (**A**) or 10 µg (**B**) of the indicated mAb delivered intraperitoneally. The median baseline-corrected weight of each treatment group is plotted. Error bars represent the 95% confidence interval. Number of animals is reported in legend.

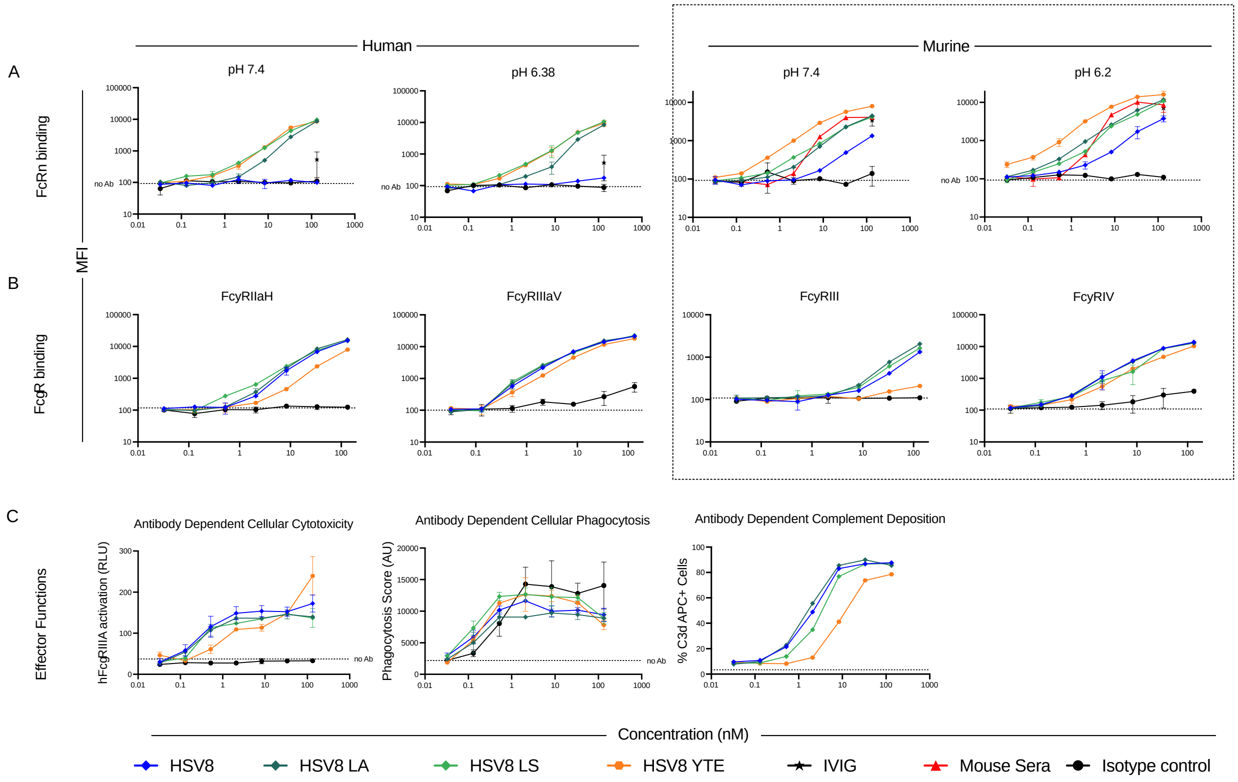


**Figure S2 (related to Figures 1 and 2)**: **HSV8 binding to FcRn and FcγR and effector functions. A.** FcRn binding profiles for HSV8 variants bound to gD-2 and detected with human (left) or mouse (right) FcRn tetramerized with streptavidin-PE at indicated pH. **B.** FcγR binding profiles for HSV8 variants to human (left) or mouse (right) FcγR. **C**. Ability for HSV8 variants to mediate ADCC (left), ADCP (center), or complement deposition (right).

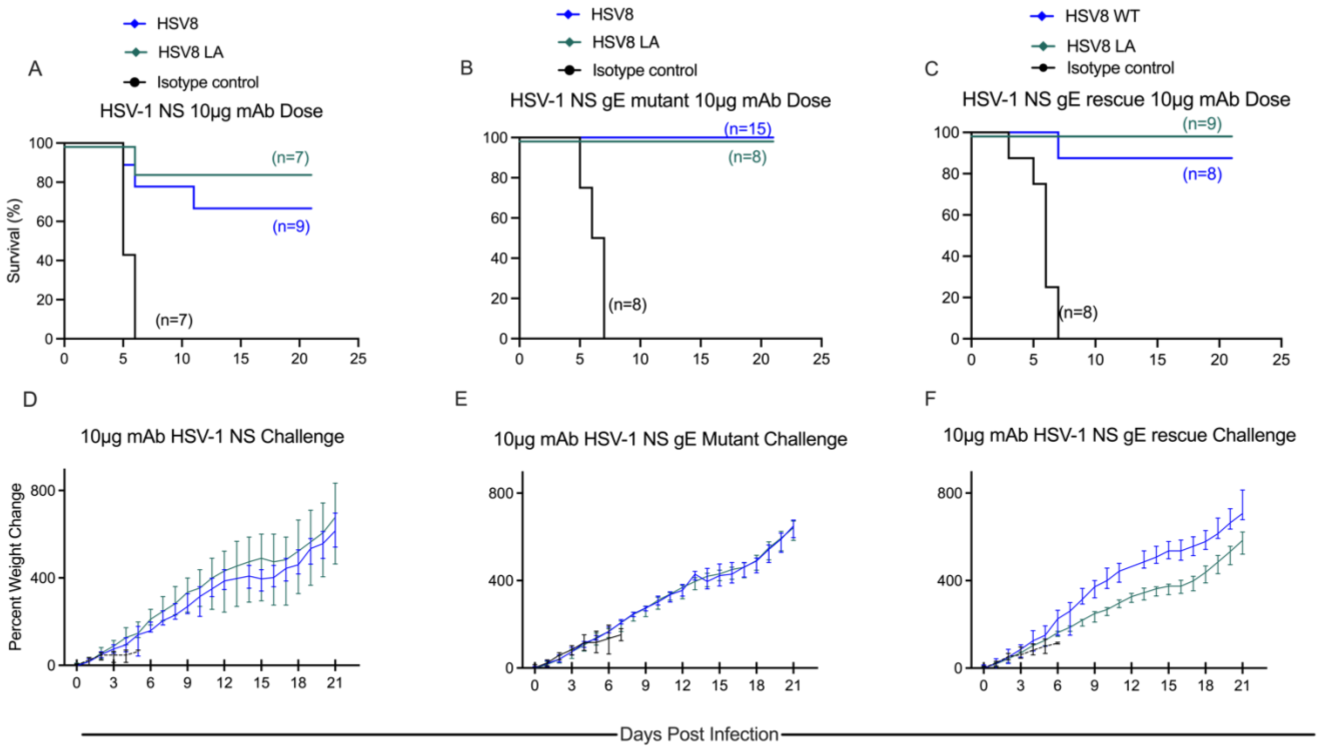


**Figure S3 (related to Figure 6): A 10 µg dose of HSV8 variants sufficiently protects mice from gE-modulated viral challenge.** mAbs (10 µg) were delivered intraperitonially to 2-day old pups immediately before a lethal ($1x10^4$ PFU) challenge with HSV-1 NS (**A**), HSV-1 NS gE mutant (**B**), or HSV-1 NS gE rescue (**C**). Weight gain of pups following lethal challenge with HSV-1 NS (**D**), HSV-1 NS gE mutant (**E**), or HSV-1 NS gE rescue (**F**).

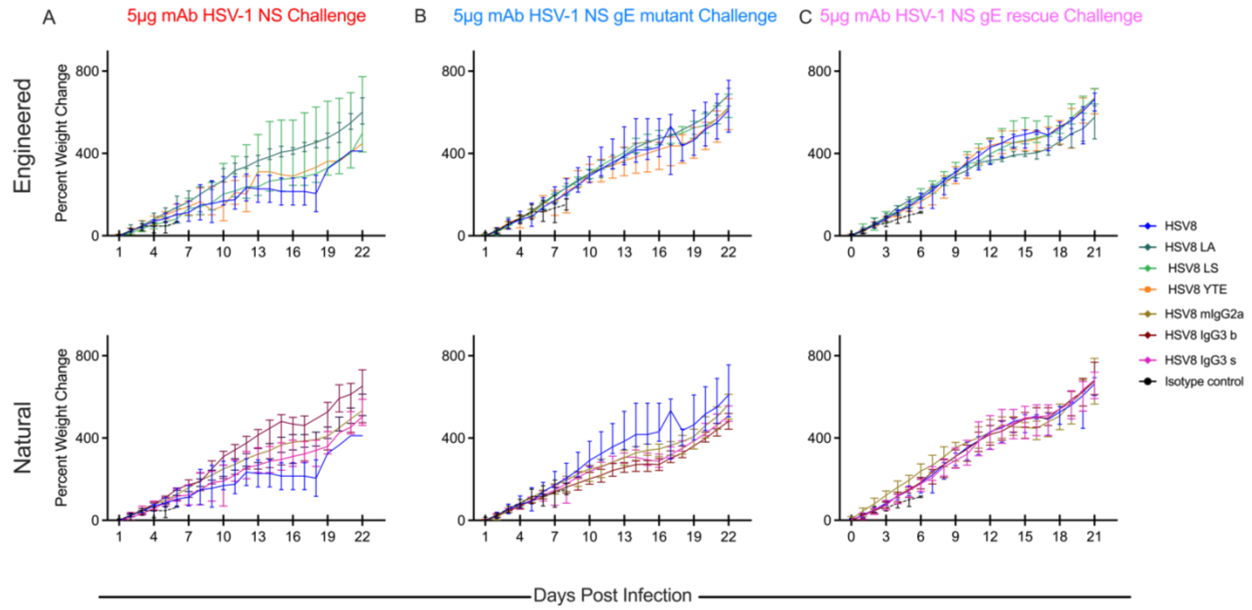


**Figure S4 (related to Figure 6): Percent weight change of mice receiving engineered or natural HSV8 variants.** mAbs (5 μg) were delivered intraperitonially to 2-day old pups immediately before a lethal (1x10$^4$ PFU) challenge with HSV-1 NS (**A**), HSV-1 NS gE mutant (**B**), or HSV-1 NS gE rescue (**C**)**.** Percent weight gain of pups following lethal challenge with indicated HSV-1 NS strains**.**

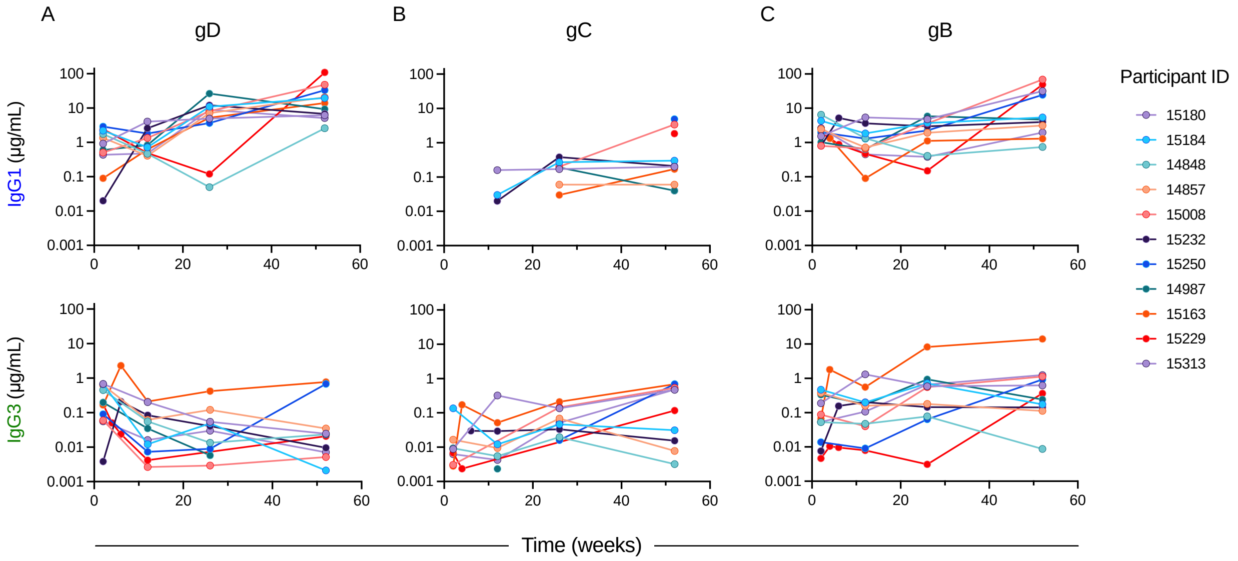


**Figure S5 (related to Figure 7): Longitudinal antibody profiling of individuals with primary HSV-1 infection.** Levels of IgG1 (top) and IgG3 (bottom) antibodies against HSV-1 gD (**A**), gC (**B**) or gB (**C**) over time.
